# Acquired fluoroquinolone resistance genes in corneal isolates of *Pseudomonas aeruginosa*

**DOI:** 10.1101/2020.05.17.100396

**Authors:** Mahjabeen Khan, Stephen Summers, Scott A Rice, Fiona Stapleton, Mark D P Willcox, Dinesh Subedi

## Abstract

Fluroquinolones are widely used as an empirical therapy for pseudomonal ocular infections. Based on increasing reports on acquired fluroquinolone resistance genes in clinical isolates of *Pseudomonas aeruginosa*, we investigated 33 strains of *P. aeruginosa* isolated from the cornea of microbial keratitis patients in India and Australia between 1992 and 2018 to understand the prevalence of acquired fluroquinolone resistance genes in ocular isolates and to assess whether the possession of those genes was associated with fluoroquinolone susceptibility. We obtained the whole genome sequence of 33 isolates using Illumina MiSeq platform and investigated the prevalence of two fluoroquinolone resistance genes *crpP* and *qnrVC1*. To examine the associated mobile genetic elements of *qnrVC1* positive strains, we obtained long read sequences using Oxford Nanopore MinION and performed hybrid assembly to combine long reads with Illumina short sequence reads. We further assessed mutations in QRDRs and antibiotic susceptibilities to ciprofloxacin, levofloxacin and moxifloxacin to examine the association between resistance genes and phenotype. Twenty strains possessed *crpP* in genetic islands characterised by possession of integrative conjugative elements. The *qnrVC1* gene was carried by four isolates on class I integrons and Tn*3* transposons along with aminoglycoside and beta-lactam resistance genes. We did not observe any evidence of plasmids carrying fluroquinolone resistance genes. Resistance to fluroquinolones was observed in those strains which possessed *crpP, qnrVC1* and that had QRDRs mutations. The presence of *crpP* was not a sole cause of fluroquinolone resistance.

## Introduction

*Pseudomonas aeruginosa* is a highly adaptable opportunistic pathogen which is ubiquitously present in the environment. This bacterium is naturally resistant to many antimicrobials and can acquire antibiotic resistance through mutations in chromosomal genes and lateral gene transfer [1, 2]. *P. aeruginosa* is associated with different types of human infections and because of emerging multidrug-resistant strains, these infections are major global public health concerns [3].

Fluoroquinolones are broad spectrum and widely prescribed antibiotics to treat pseudomonal infections including ocular infections [4–6]. Fluoroquinolone resistance in various clinical isolates is on the rise [7]. For example, a single centre study has shown that the prevalence of fluoroquinolone resistance *P. aeruginosa* increased from 15% to 41% in ten years [8]. This increase in fluoroquinolone resistance has been linked to the excessive use of the antibiotics

[9]. The rate of isolation of fluoroquinolone resistant strains also depends on the type of infections; nosocomial isolates are more resistant than isolates from non-nosocomial sources

[10]. In general, fluoroquinolone resistance is relatively low in ocular isolates of *P. aeruginosa* compared to other infections [11]. However, higher resistance rates have been reported in ocular isolates in certain regions of the world and like other systemic infections this rate has been increasing over time [12]. This has raised the concerned that the horizontal transfer of fluoroquinolone resistance genes can be associated with spread of fluoroquinolone resistance in ocular isolates.

Mutations that alter target sites (DNA gyrase [*gyrA/gyrB*] and topoisomerase IV [*parC/parE*]) and increased membrane permeability are common mechanisms of fluoroquinolone resistance in *P. aeruginosa* [1, 13–15]. In other Gram-negative bacteria, fluoroquinolone resistance genes such as *qnr* have been shown to be carried on plasmids [16, 17]. The gene encodes a pentapeptide repeat protein which protects DNA gyrase and topoisomerase from the action of fluroquinolones [18]. In contrast, carriage of *qnr* on plasmid is very rare in *P. aeruginosa* [19]. Despite this low carriage, recent studies have identified fluoroquinolone resistance genes in certain mobile genetic elements. *CrpP* and *qnrVC* can be carried on the *P. aeruginosa* mega plasmids pUM505 and pBM413, respectively [20, 21]. In addition, *qnrVC* was found to be associated with a class I integron, which also carried beta-lactamase genes [20, 22]. These studies were based on a single strain and the prevalence of acquired fluoroquinolone resistance genes in *P. aeruginosa* remains unclear. This led us to examine mobile fluoroquinolone resistance genes in *P. aeruginosa*. Given the concern of increasing fluoroquinolone resistance in ocular isolates, we have undertaken sequence analysis of 33 ocular isolates of *P. aeruginosa*, isolated from corneal ulcers in the last 25 years to assess whether the possession of fluoroquinolone resistance genes was associated with fluoroquinolone susceptibility and whether resistance genes carrying strains had any genetic commonalities.

## Materials and Methods

### *Pseudomonas aeruginosa* strains

Isolates in this study were collected from the cornea of microbial keratitis patients in India and Australia between 1992 and 2018. They comprise 20 isolates collected for this study and used for the antibiotic susceptibility testing and whole genome sequencing. Information on antibiotic susceptibility and the whole genome data of 13 isolates was obtained from our previous study [23] (Supplementary Table 1). All isolates were retrieved from the culture collection of School of Optometry and Vision Science, the University of New South Wales, Australia without identifiable patients’ data.

### Antibiotic susceptibility testing of *Pseudomonas aeruginosa* strains

Susceptibility of *P. aeruginosa* isolates to ciprofloxacin (Sigma-Aldrich, Inc., St. Louis Missouri, USA), levofloxacin (Sigma-Aldrich, Inc.,) and moxifloxacin (European Pharmacopoeia, Strasbourg, Cedex France) was investigated using the broth microdilution method following the protocol of Clinical and Laboratory Standard Institute [24] [25]. The MIC was taken as the lowest concentration of an antibiotic in which no noticeable growth (turbidity) was observed and the break point was established according to published standards [26]. Based on MICs, the isolates were categorised into four groups, susceptible (≤ resistance break point), resistant (> resistance break point – 32 μg/mL), highly resistant (> 32 – 128 μg/mL) and very highly resistant (>128 μg/mL) for the analysis.

### DNA extraction and Illumina sequencing

Bacterial DNA was extracted from overnight cultures using a DNeasy Blood and Tissue Kit (Qiagen, Hilden, Germany) following the manufacturer’s instructions. The extracted DNA was quantified and purity-checked using Nanodrop (NanoDrop Technologies, Wilmington, DE, USA), Qubit fluorometer (Life Technologies, Carlsbad, CA, USA), and 1% agarose gel electrophoresis. The extracted DNA was dried for transport to the sequencing facility at Singapore Centre for Environmental Life Sciences Engineering, Singapore. DNA was sequenced on MiSeq (Illumina, San Diego, CA, USA) generating 300 bp paired end reads. The paired-end library was prepared using Nextera XT DNA library preparation kit (Illumina, San Diego, CA, USA). All the libraries were multiplexed on one MiSeq run.

### Bioinformatics analysis of short read sequences and construction of phylogenies

The quality of raw reads was analysed using online tool FastQC v0.117 (https://www.bioinformatics.babraham.ac.uk/projects/fastqc). Adaptor sequences were removed using Trimmomatic v0.38 with quality and length filtering (SLIDINGWINDOW:4: 15 MINLEN:36) [27]. The reads were de-novo assembled using Spades v3.13.0 with the programs’ default setting [28] followed by annotation using Prokka v1.12 [29]. To examine acquired resistance genes, the genomes were uploaded into online database ResFinder v3.1 of Centre for Genomic Epidemiology, DTU, Denmark [30]. The contigs carrying *crpP* and *qnrVC1* genes were selected and analysed for mobile genetic elements using BLAST, Integron finder v1.5.1 [31] and IS finder [32]. The genome was visualised and manually modified using Geneious prime v2019.2 [33]. A figure of BLAST comparison was generated using EasyFig v2.2.2 [34]. To identify mutations in the QRDRs (*gyrA, gyrB, parC* and*parE*), the genome sequences were analysed using Snippy v4.2 with program’s default settings (https://github.com/tseemann/snippy). The genomes were also examined for the presence of type IV secretion factors (*exoU* and *exoS*) using BLAST search, and for the presence of the CRISPR cas system using the CRISPRcasFinder online tool [35]. Codon adaptation index (CAI) [36] was examined to understand the possible expression of different orthologues of *crpP* using CAIcal [37, 38]. 50s ribosomal protein L19 (*rplS*), which is a highly expressed chromosomal gene was included as the reference to show the difference in expression level between chromosomal gene and acquired genes.

Core genome single nucleotide polymorphisms (SNPs) were identified using Parsnp v from Harvest Suite [39] using settings to exclude SNPs identified in regions that had arisen by recombination. The core genome SNPs were used to construct a maximum likelihood phylogenetic tree. All genomes were examined for multi-locus sequence type (MLST) using the MLST database. Nucleotide sequences of all MLST locus were extracted and concatenated to use in Bayesian phylogenetic analyses using BEAST2 v2.4.7 with the following parameters: gamma site heterogeneity model, Hasegawa-Kishino-Yano (HKY) substitution model and relaxed-clock log-normal [40]. BEAST 2 output was summarized using TreeAnnotator with a 5% burn in. The phylogenetic trees were visualised using iTol v4 [41].

### MinION sequencing and analysis

Wizard Promega DNA extraction kits (Promega, Madison, WI) was used to extract DNA from overnight broth culture. DNA was quantified and transported as mentioned above. Long reads libraries were prepared using a rapid sequencing kit (RAD-SQK004, ONT, Oxford, UK) and subsequent sequencing was conducted using the MinIon flowcell (R9.4.1) for 48 hours. Long reads were basecalled using Guppy v3.3.0 and adapters removed using Porechop v0.2.4. Assembly of both long and short reads into a hybrid genome assembly was achieved with Unicycler v0.4.3 [42], opting for default parameters and using only short reads with merging pairs. Following assembly, all assemblies were assessed for quality using Quast v5.0.2.

## Results

### Population structure, phylogeny and fluoroquinolone resistance of *P. aeruginosa* strains

Whole genomes were analysed from 33 corneal isolates of *P. aeruginosa*, 20 of which were sequenced as a part of this study and 13 genome that were published previously [23]. All strains were isolated between 1992 and 2018 in India (19 isolates) or Australia (14 isolates) (Supplementary Table 1). Draft genomes were mapped against the reference genome *P. aeruginosa* PAO1 and a total of 202,232 SNPs were observed among the 33 isolates, which were used to construct core genome phylogeny using Parsnp v1.2 [43]. Similar to previous reports [23, 36, 44, 45], the phylogenetic tree based on the core genome revealed two major clades (Fig 1). These clades followed a similar pattern as for other previously described strains of *P. aeruginosa*, where *exoU* carrying strains clustered together in a single clade (Phylogroup 2) [23]. Our results showed that 18 (of 33) isolates carried *exoU*, of which 16 were clustered together in phylogroup 2, which contained predominantly Indian isolates. The detection rate of *exoU* was 68% in Indian isolates and 36% in Australian isolates.

**Figure 1.**
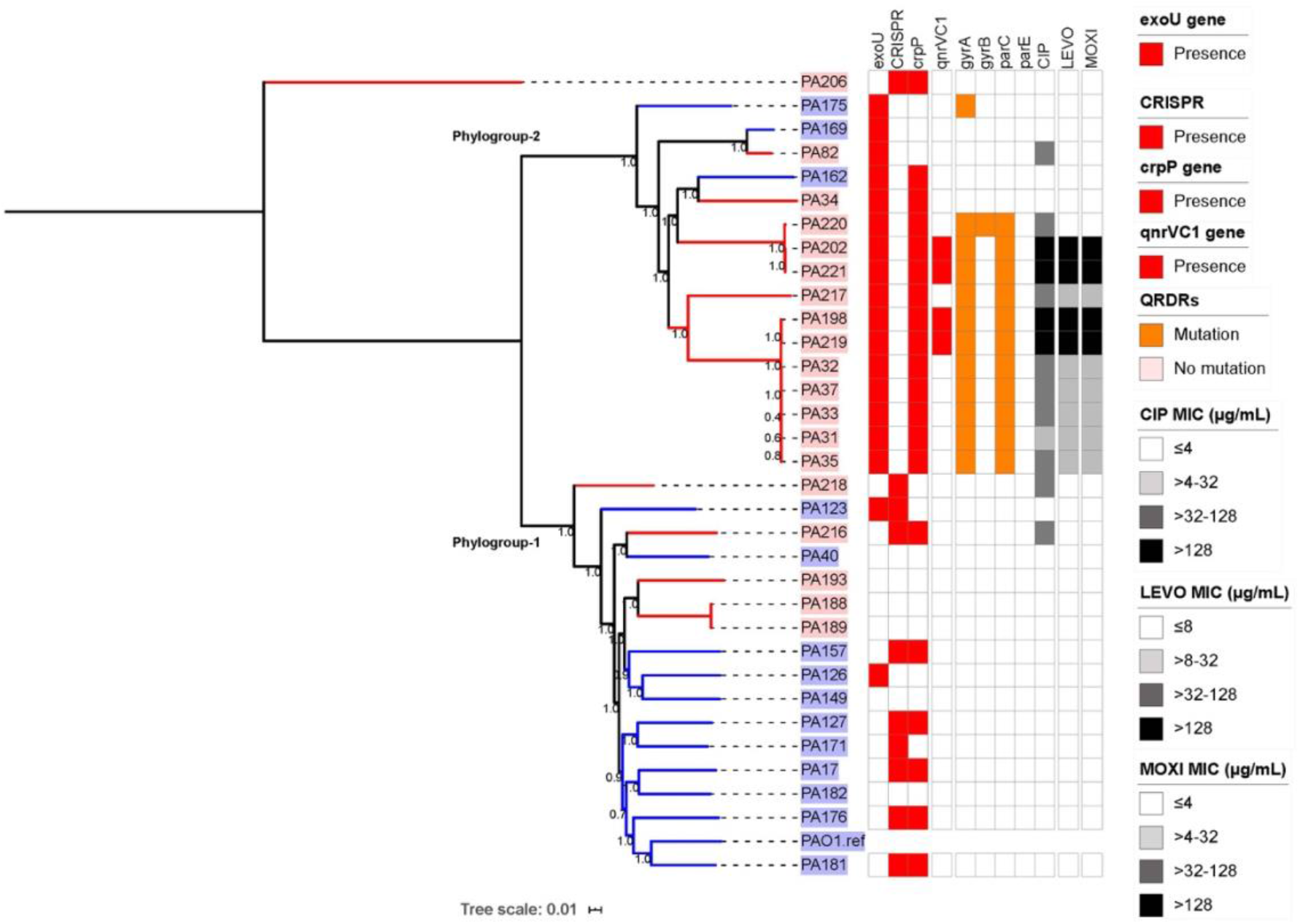
Maximum likelihood phylogenetic tree based on core genome SNPs analysis using *Pseudomonas aeruginosa* PAO1 as the reference, excluding SNPs identified in regions that had arisen by recombination, using the default parameters of Parsnp v1.2 [43]. Isolates from India are labelled red and Australian isolates are labelled blue. Numbers given at the nodes represent bootstrap values. The presence of *crpP, exoU, qnrVC1*, and CRISPR cas are represented by red squares. Orange squares represent the presence of mutations in the quinolone resistance determining regions (QRDRs) and fluoroquinolone (CIP = Ciprofloxacin; LEVO = Levofloxacin; and MOX = Moxifloxacin) susceptibilities are shown as a heat map with the ranges indicated in the figure. The figure was drawn using iTol v4 [41].

We further examined acquired fluoroquinolone resistance genes, mutations in quinolone resistance determining region (QRDRs), the presence of CRISPR genes and susceptibility to three different fluoroquinolones (Fig 1). None of the Australian isolates regardless of phylogenic grouping were resistant to fluoroquinolones. In contrast, 75% of isolates of phylogroup 2 were resistant to at least one fluoroquinolone. Of the 33 strains, 73.7 % (14 out of 19) of Indian and 42.8% (6 out of 14) of Australian isolates possessed *crpP*, which has been recently shown to be on a plasmid (pUM505) and associated with ciprofloxacin resistance [20]. However, eight (40%) *crpP* carrying strains in the current study regardless of country of isolation were not resistant to the fluoroquinolones, including ciprofloxacin (Fig 1). Eleven out of 14 fluoroquinolone resistance strains had mutations in both *gyrA* and *parC* and all except one carried the *exoU* gene. In addition, four strains from the latter cohort of 11 strains carried another fluoroquinolone resistance gene, *qnrVC1*, in combination with mutations in *gyrA* and *parC*, and this was associated with a very high MIC (>128 μg/mL) to all three fluoroquinolones (Fig 1).

Given that *exoU, crpP* and *qnrVC1* are components of the accessory genome which is mostly shaped by the CRISPR-Cas system, a bacterial defence system against foreign DNA [46], we searched isolates for CRISPR-Cas genes using the CRISPRCasFinder database [35] following software default parameters. Only one of the *exoU*, and none of the *qnrVC1* carrying strains, were positive for CRISPR-Cas genes. However, CRISPR-Cas was observed in seven *crpP* positive strains, which did not possess *exoU* and/or *qnrVC1* (Fig 1).

### Bayesian phylogenetic reconstruction

To examine the evolutionary trends of isolates, a Bayesian analysis was performed based on MLST sequences. The evolutionary tree was constructed by BEAST, which uses a molecular clock model to estimate time-measured phylogenies [40]. The result suggested that the most recent common ancestor of isolates of this study appeared approximately 50 years ago and the branching of fluoroquinolone resistant strains (except PA82), which had mutations in QRDRs and/or had acquired resistance *qnrVC1* within each subclade, occurred between 1988 and 2000 (Fig 2). This coincided with the time when ciprofloxacin and ofloxacin were introduced as antibiotic treatments [47]. The analysis suggests that acquisition of *crpP* and *exoU* occurred at least 50 years ago.

**Figure 2.**
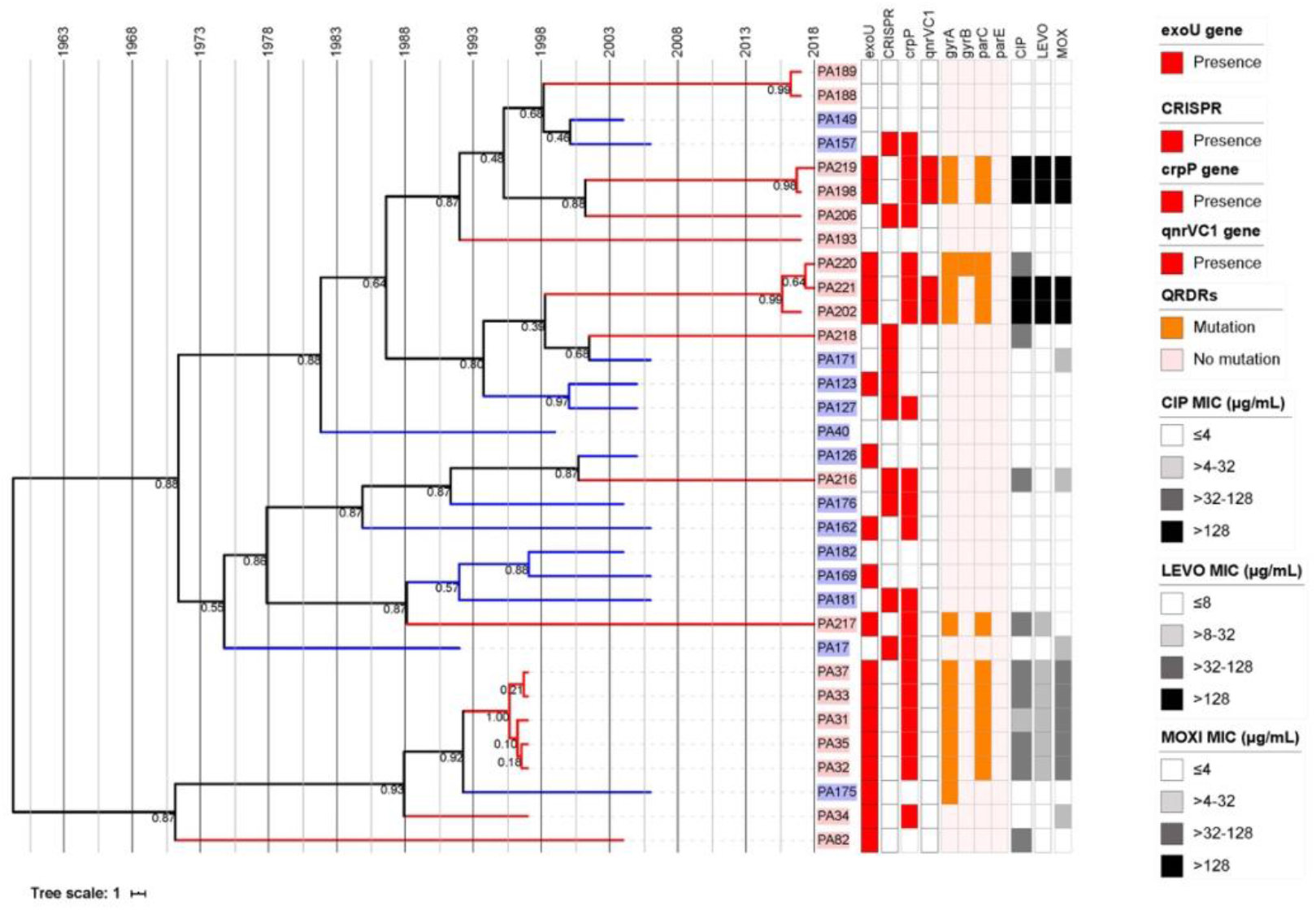
Consensus tree of 33 *P. aeruginosa* isolates, based on Bayesian evolutionary analysis by sampling trees (BEAST) of concatenated multi-locus sequence type (MLST) under strict clock analysis [40]. The tip of the tree was constrained by date of isolation. The time scale is shown in years at the top and each internal node is labelled with posterior probability limit. Isolates from India are labelled red and Australian isolates are labelled blue. The presence of genes *crpP, exoU, qnrVC1*, and CRISPRcas are represented by red squares. Orange square represents presence of mutations in quinolone resistance determining region (QRDRs), Fluoroquinolone (CIP = Ciprofloxacin; LEVO = Levofloxacin; and MOX = Moxifloxacin) susceptibilities are shown as heat maps in the grey scale indicated in the figure. The figure was drawn using iTol v4 [41].

### Prevalence of the *crpP* gene

A recent study has shown that *crpP* encodes for a protein that is associated with increased resistance to ciprofloxacin [20]. In the current study, 63% of isolates possessed this gene, however nine out of the 20 *crpP* carrying strains were not resistant to ciprofloxacin (Fig 1). Single nucleotide polymorphisms in the *crpP* sequences from all strains were compared to the sequence of *crpP* in pUM505 using local BLASTn. The BLAST matrix (Table 1) showed that the nucleotide sequences of *crpP* genes varied between 100% and 93.43%, indicating the possibility of differences in the activity of the encoded protein. Codon usage, presented as codon adaptation index [36] (Fig 3) was examined to understand the possible expression of different orthologues of *crpP* [38]. The 50s ribosomal protein L19 (*rplS*), which is a highly expressed chromosomal gene was included in this analysis to show the difference in expression between chromosomal and acquired genes. A CAI between 0.36 and 0.39 was observed in different orthologues of *crpP*, which while low, corresponded to divergence of sequences. However, the sequence divergence was not associated with resistance to fluoroquinolones. For example, the nucleotide sequence of *crpP* of a resistant strain PA32 and a sensitive strain PA127 were 98.48% identical to that of pUM505.

This study also investigated the presence of the *crpP* homologues in NCBI databases using local BLAST with a cut off of 80% coverage and 90% sequence identity to examine the distribution of *crpP* gene in the bacterial database including plasmids as of 2019-06-15. The BLAST search against *P. aeruginosa* complete genome databases returned 52% matches and this rate of detection is broadly comparable to *crpP* prevalence observed in the current strains. No BLAST matches other than *P. aeruginosa* were observed with this search parameter. Within the *P. aeruginosa* plasmid database, two matches were observed against 45 complete plasmids using the parameters mentioned above. A CrpP-like protein, which has sequence identity of 10-46% with *P. aeruginosa crpP*, has been observed in species of the *Enterobacteriaceae* [48]. This result suggested that *crpP* can be transferred between species or strains.

**Table 1:** BLAST matrix showing nucleotide identity (in percentage) of *crpP* sequences of each isolate. Colour intensity corresponds to percentage identity.

|  | pUM505* | PA31 | PA32 | PA33 | PA35 | PA37 | PA127 | PA162 | PA220 | PA17 | PA34 | PA157 | PA206 | PA217 | PA181 | PA216 | PA202 | PA219 | PA221 | PA176 | PA198 |
| --- | --- | --- | --- | --- | --- | --- | --- | --- | --- | --- | --- | --- | --- | --- | --- | --- | --- | --- | --- | --- | --- |
| pUM505* |  | 98.48 | 98.48 | 98.48 | 98.48 | 98.48 | 98.48 | 98.48 | 98.48 | 97.98 | 97.98 | 97.98 | 97.98 | 97.98 | 96.46 | 95.45 | 94.95 | 94.95 | 94.95 | 94.44 | 94.44 |
| PA31 | 98.48 |  | 100 | 100 | 100 | 100 | 100 | 98.99 | 100 | 99.49 | 98.48 | 98.48 | 98.48 | 98.48 | 97.98 | 95.96 | 95.45 | 95.45 | 95.45 | 94.95 | 94.95 |
| PA32 | 98.48 | 100 |  | 100 | 100 | 100 | 100 | 98.99 | 100 | 99.49 | 98.48 | 98.48 | 98.48 | 98.48 | 97.98 | 95.96 | 95.45 | 95.45 | 95.45 | 94.95 | 94.95 |
| PA33 | 98.48 | 100 | 100 |  | 100 | 100 | 100 | 98.99 | 100 | 99.49 | 98.48 | 98.48 | 98.48 | 98.48 | 97.98 | 95.96 | 95.45 | 95.45 | 95.45 | 94.95 | 94.95 |
| PA35 | 98.48 | 100 | 100 | 100 |  | 100 | 100 | 98.99 | 100 | 99.49 | 98.48 | 98.48 | 98.48 | 98.48 | 97.98 | 95.96 | 95.45 | 95.45 | 95.45 | 94.95 | 94.95 |
| PA37 | 98.48 | 100 | 100 | 100 | 100 |  | 100 | 98.99 | 100 | 99.49 | 98.48 | 98.48 | 98.48 | 98.48 | 97.98 | 95.96 | 95.45 | 95.45 | 95.45 | 94.95 | 94.95 |
| PA127 | 98.48 | 100 | 100 | 100 | 100 | 100 |  | 98.99 | 100 | 99.49 | 98.48 | 98.48 | 98.48 | 98.48 | 97.98 | 95.96 | 95.45 | 95.45 | 95.45 | 94.95 | 94.95 |
| PA162 | 98.48 | 98.99 | 98.99 | 98.99 | 98.99 | 98.99 | 98.99 |  | 98.99 | 98.48 | 98.48 | 99.49 | 99.49 | 99.49 | 97.47 | 94.95 | 94.44 | 94.44 | 94.44 | 94.95 | 94.95 |
| PA220 | 98.48 | 100 | 100 | 100 | 100 | 100 | 100 | 98.99 |  | 99.49 | 98.48 | 98.48 | 98.48 | 98.48 | 97.98 | 95.96 | 95.45 | 95.45 | 95.45 | 94.95 | 94.95 |
| PA17 | 97.98 | 99.49 | 99.49 | 99.49 | 99.49 | 99.49 | 99.49 | 98.48 | 99.49 |  | 98.99 | 97.98 | 97.98 | 97.98 | 97.47 | 96.46 | 95.96 | 95.96 | 95.96 | 94.44 | 94.44 |
| PA34 | 97.98 | 98.48 | 98.48 | 98.48 | 98.48 | 98.48 | 98.48 | 98.48 | 98.48 | 98.99 |  | 98.99 | 98.99 | 98.99 | 97.47 | 95.45 | 94.95 | 94.95 | 94.95 | 93.43 | 93.43 |
| PA157 | 97.98 | 98.48 | 98.48 | 98.48 | 98.48 | 98.48 | 98.48 | 99.49 | 98.48 | 97.98 | 98.99 |  | 100 | 100 | 97.98 | 94.44 | 93.94 | 93.94 | 93.94 | 94.44 | 94.44 |
| PA206 | 97.98 | 98.48 | 98.48 | 98.48 | 98.48 | 98.48 | 98.48 | 99.49 | 98.48 | 97.98 | 98.99 | 100 |  | 100 | 97.98 | 94.44 | 93.94 | 93.94 | 93.94 | 94.44 | 94.44 |
| PA217 | 97.98 | 98.48 | 98.48 | 98.48 | 98.48 | 98.48 | 98.48 | 99.49 | 98.48 | 97.98 | 98.99 | 100 | 100 |  | 97.98 | 94.44 | 93.94 | 93.94 | 93.94 | 94.44 | 94.44 |
| PA181 | 96.46 | 97.98 | 97.98 | 97.98 | 97.98 | 97.98 | 97.98 | 97.47 | 97.98 | 97.47 | 97.47 | 97.98 | 97.98 | 97.98 |  | 93.94 | 93.43 | 93.43 | 93.43 | 93.43 | 93.43 |
| PA216 | 95.45 | 95.96 | 95.96 | 95.96 | 95.96 | 95.96 | 95.96 | 94.95 | 95.96 | 96.46 | 95.45 | 94.44 | 94.44 | 94.44 | 93.94 |  | 99.49 | 99.49 | 99.49 | 93.94 | 93.94 |
| PA202 | 94.95 | 95.45 | 95.45 | 95.45 | 95.45 | 95.45 | 95.45 | 94.44 | 95.45 | 95.96 | 94.95 | 93.94 | 93.94 | 93.94 | 93.43 | 99.49 |  | 100 | 100 | 93.43 | 93.43 |
| PA219 | 94.95 | 95.45 | 95.45 | 95.45 | 95.45 | 95.45 | 95.45 | 94.44 | 95.45 | 95.96 | 94.95 | 93.94 | 93.94 | 93.94 | 93.43 | 99.49 | 100 |  | 100 | 93.43 | 93.43 |
| PA221 | 94.95 | 95.45 | 95.45 | 95.45 | 95.45 | 95.45 | 95.45 | 94.44 | 95.45 | 95.96 | 94.95 | 93.94 | 93.94 | 93.94 | 93.43 | 99.49 | 100 | 100 |  | 93.43 | 93.43 |
| PA176 | 94.44 | 94.95 | 94.95 | 94.95 | 94.95 | 94.95 | 94.95 | 94.95 | 94.95 | 94.44 | 93.43 | 94.44 | 94.44 | 94.44 | 93.43 | 93.94 | 93.43 | 93.43 | 93.43 |  | 100 |
| PA198 | 94.44 | 94.95 | 94.95 | 94.95 | 94.95 | 94.95 | 94.95 | 94.95 | 94.95 | 94.44 | 93.43 | 94.44 | 94.44 | 94.44 | 93.43 | 93.94 | 93.43 | 93.43 | 93.43 | 100 |  |

**Figure 3.**
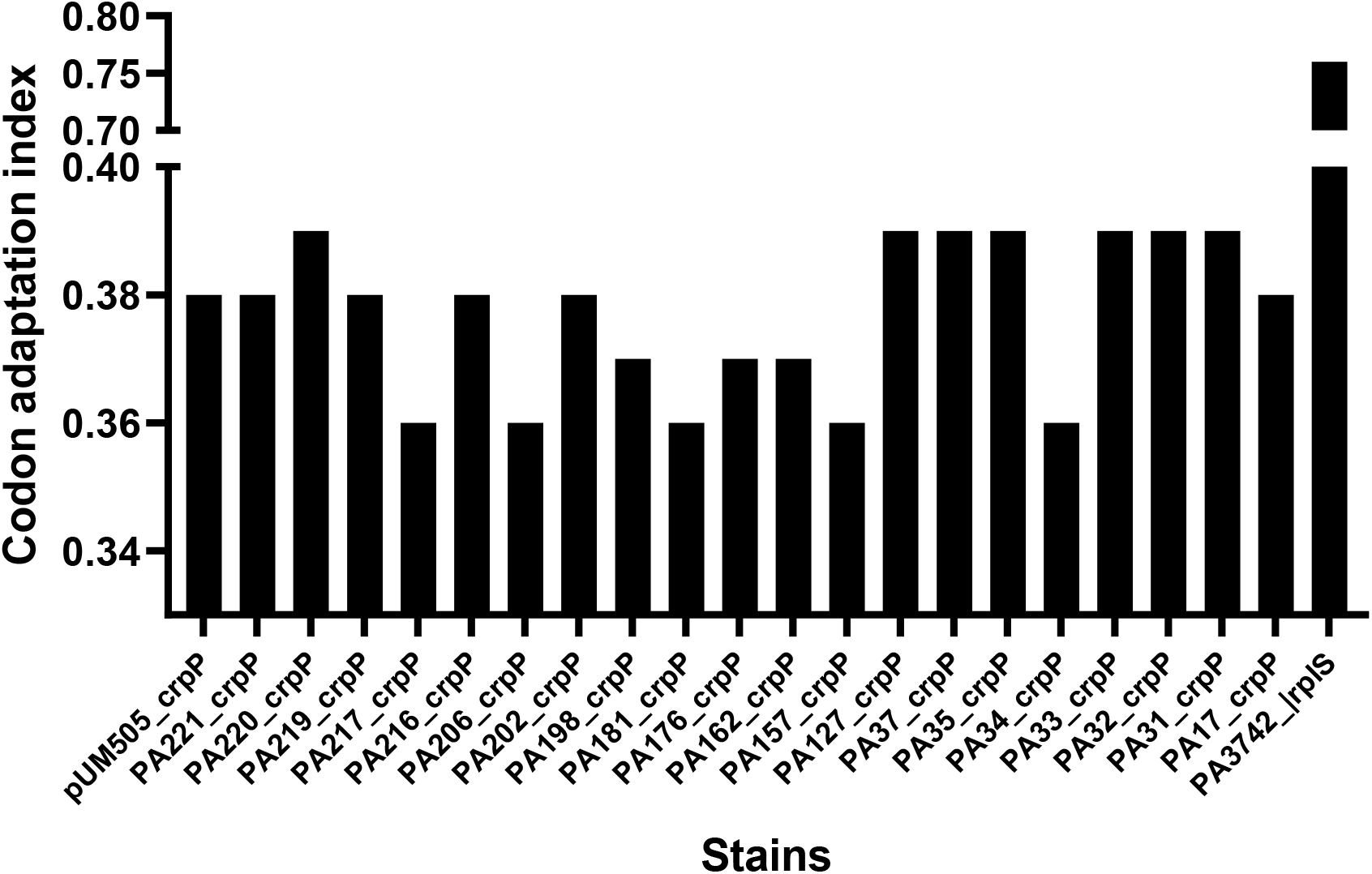
Codon adaptation index of *crpP* orthologues of *P. aeruginosa* strains. PA3742_rplS denotes 50s ribosomal protein L19 (rplS) of *P. aeruginosa* PAO1.

### *crpP* associated mobile genetic elements

Although the *crpP* gene was first reported in plasmid pUM505 of a clinical strain of *P. aeruginosa* [20], BLAST analysis, as mentioned above, revealed this gene was mostly associated with the chromosome and its presence in plasmids was rare. The *crpP* containing contigs of each isolate were assessed to determine similarities between associated genomic islands. The *crpP* flanking region of pUM505 is homologous to the pathogenicity island PAPI-1 of *P. aeruginosa* PA14 [49] and pKLC102 of *P. aeruginosa* C [50, 51], which is an integrative conjugative element and is associated with carriage of virulence genes. A PAPI-1-like genomic island has also been reported in a complete genome of the ocular isolate PA34 [52]. Using PAPI-1 of PA34 as a reference, the contigs of each strain were compared and this showed that there were similarities between contigs. Those common regions were extracted and annotated manually using Geneious Prime v2019.1.3 [33]. After annotation, all of the extracted regions were compared using Easyfig [34]. The region that has been shared amongst all isolates is presented in Fig 4, where the colour code represents open reading frames. While the size of *crpP* carrying genomic islands differed between strains, the exact size of islands could not be ascertained because of the use of draft genomes for analysis. Nevertheless, three genes (integrative conjugative element, an unknown protein and single-strand binding protein) upstream and one gene (type I DNA topoisomerase) downstream of *crpP* were common in all isolates and this indicates that these genes may be involved in the transfer of *crpP* gene. A characteristic type IV pili synthesis operon was associated with 75% of the *crpP* positive strains. There was no association between size of *crpP* islands and fluoroquinolone resistance.

**Figure 4.**
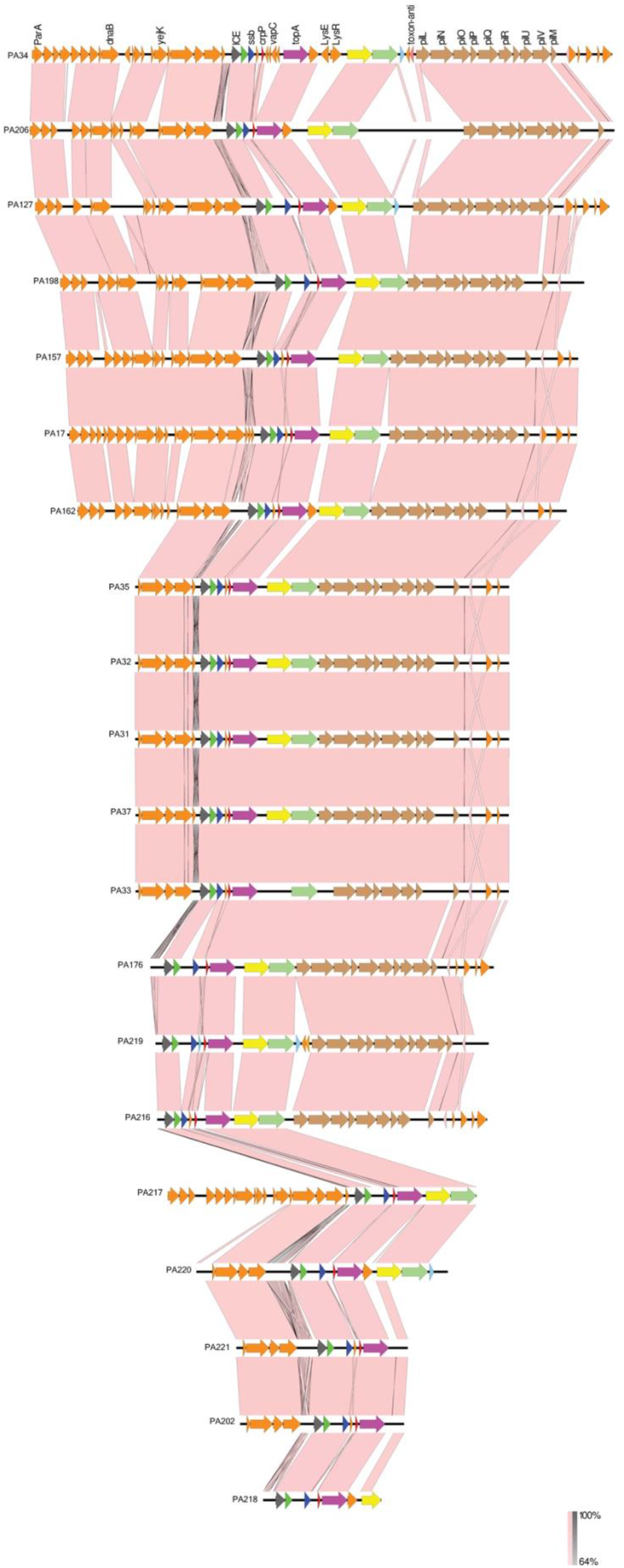
Comparison of crpP-associated genomic islands of *P. aeruginosa* strains. Protein-coding regions are represented by the orange arrows and common key features/associated genes among all strains are shown in various coloured arrows. The gradient pink and grey shading represent regions of nucleotide sequence identity (100% to 64%) in forward and reverse directions respectively, determined by BLASTn analysis. Figures are drawn to scale using Easyfig [34]. (parA= plasmid partition protein A, dnaB= replicative DNA helicase, yejK= nucleoid-associated protein, ICE= integrating conjugative element protein, ssb= single-stranded DNA-binding protein, crpP= ciprofloxacin resistance protein, vapC= tRNA(fMet)-specific endonuclease, topA= type I DNA topoisomerase, LysE= LysE family translocator, LysR= LysR family transcriptional regulator, toxin-anti= type II toxin-antitoxin system RelE/ParE family toxin, pilL-pilM= type IV pilus synthesis operon).

### Prevalence of quinolone resistance protein (*qnr* gene)

The *qnr* is commonly carried on plasmids, especially in the *Enterobacteriaceae*. A transferable variant of the *qnr* gene, *qnrVC1* was first identified in a class I integron in *Vibrio cholerae* 01 [53]. Several *qnrVC* alleles have also been reported to be carried in various bacteria including *P. aeruginosa* [18, 54]. For example, a recent study had reported that 2.3% of clinical strains of *P. aeruginosa* carried *qnrVC* [55]. In the current study, 12% (four out of 33) of isolates possessed *qnrVC1*. All these four were isolated in 2017 and 2018. We further examined the prevalence of *qnrVC1* homologues in complete chromosomes and plasmids of other bacterial species in NCBI database using BLAST search. We observed that *qnrVC1-like* proteins were present in several bacterial species. Upon limiting the BLAST criteria to 100% coverage and 98% identity, the search returned with 19 matches from species of *Pseudomonas, Vibrio* and *Aeromonas*. (Supplementary Figure 1). This suggests that the *qnrVC1* gene is prevalent in various bacterial families. Curiously, the *qnrVC1* gene was not identified in the plasmid database of NCBI. Furthermore, *qnrVC1* had a G + C content of 37%, which is considerably different from that of the *P. aeruginosa* genome (66%). This evidence along with the presence of *qnrVC1* homologues in several bacterial species indicates that *qnrVC1* is a horizontally acquired gene in *P. aeruginosa*, which is potentially acquired through inter-species or inter-genera transfer.

### *qnrVC1* associated mobile genetic elements

We further examined the flanking nucleotide sequences of *qnrVC1* to understand the mobile genetic elements associated with transfer of the gene. Our initial analysis found that *qnrVC1* was present in small contigs (contig size <5000 bp) and this limited our aim to examine mobile genetic elements. The four *qnrVC1* positive strains were then analysed using the long-read sequencing technique ONT minION (ONT, Oxford). With hybrid assembly of long and short sequencing reads, we archived long contigs ranging in size from 400 Kbp to 2000 Kbp (PA202 N50 = 680,290; PA219 N50 = 2,159,014; PA221 N50 = 2,110,541) in three isolates and a complete and closed genome of one strain PA198 (N50 = 7,188,952).

The *qnrVC1* gene was present in two different types of mobile genetic elements. In strains PA198 and PA219, the gene was integrated into Tn*3* transposons, which have 99% identity with one another (Fig 5). In addition, the Tn*3* transposon carried other antibiotic resistance genes associated with aminoglycoside 3-phosphotransferase and tetracycline resistance. BLAST comparison revealed recombination in various places within the transposon, most notably in the areas flanking the *qnrVC1-VOC* and *tetR-tetA* genes (Fig 5), which represents the site of integration of these genes. The role of VOC protein in antibiotic resistance is not well known. However, evidence suggests that proteins of VOC the family include beomycin and fosfomycin resistance proteins [56]. Integration of VOC with other antibiotic resistance genes leads us to hypothesise that VOC can be associated with antibiotic resistance. Further research is required to investigate its role in antibiotic resistance in *P. aeruginosa*.

**Figure 5.**
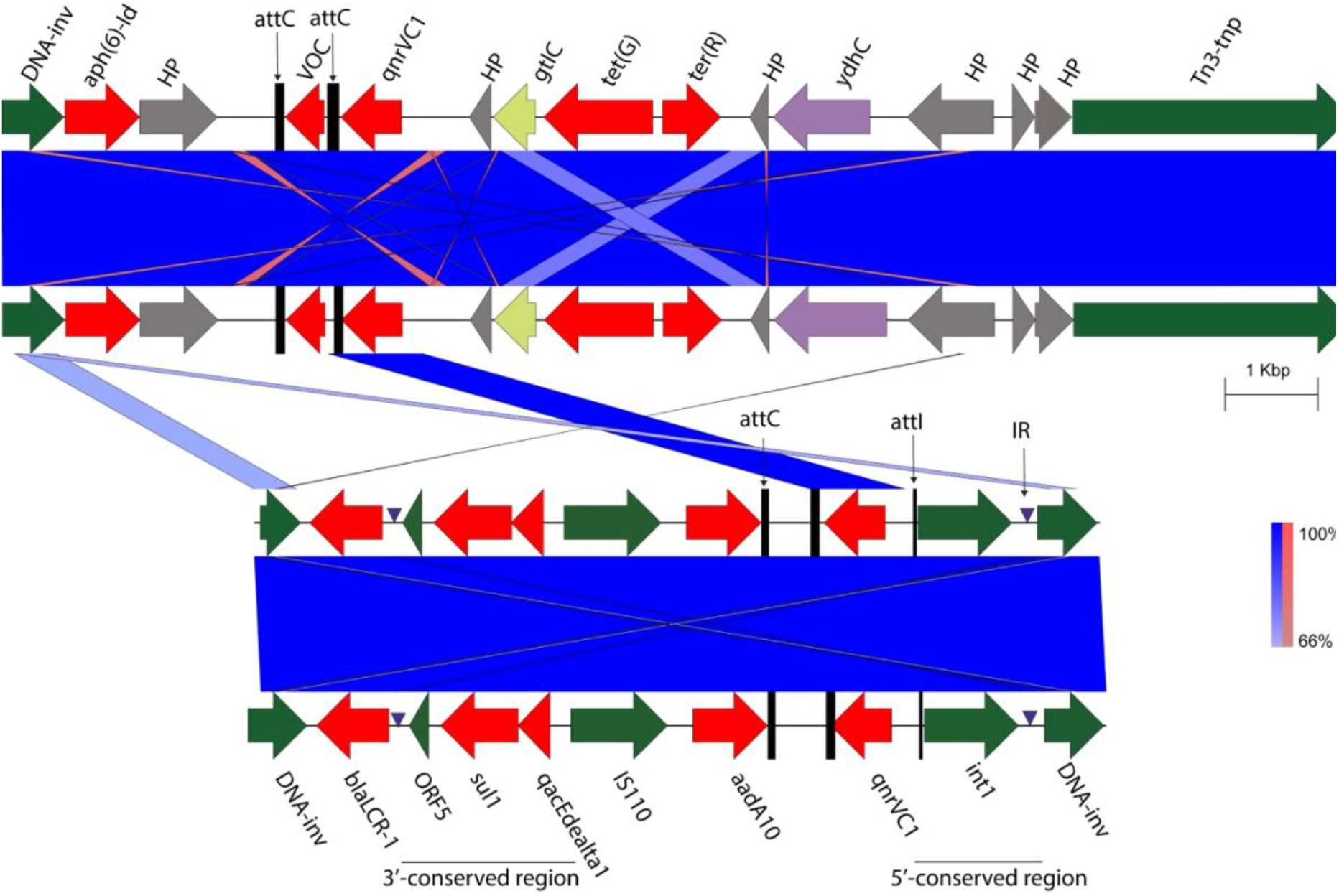
Comparison of *qnrVC1*-associated genomic islands of *P. aeruginosa* strains. Protein-coding regions are represented by the arrows and common key features/associated genes among all strains are shown in various coloured arrows. The gradient blue and red shading represent regions of nucleotide sequence identity (100% to 65%) in forward and reverse directions, respectively, determined by BLASTn analysis. The sequences are from strains top to bottom; PA198, PA219, PA202 and PA221. Figures are drawn to scale using Easyfig [34]. (Tn3-tnp = Tn3-transposase HP= hypothetical protein, ydhC= Inner membrane transport protein, tet(R)-tet(g) = tetracycline resistance genes, qnrVC1=quinolone resistance gene, VOC = VOC family protein, aph(6)-Id = aminoglycoside resistance protein, DNA-inv = DNA invertase, aadA10= aminoglycoside resistance protein, IS110= IS110 family transposase, qacEdealta1= quaternary ammonium compound-resistance protein, sul1= Dihydropteroate synthase, blaLCR-1=beta-lactamase gene, attC = recombination sites of gene cassette, attI = integron recombination site, IR=invert repeat).

In the other two strains, PA202 and PA221, *qnrVC1* was carried by identical class 1 integrons (Fig 5), along with two additional antibiotic resistance genes; the aminoglycoside gene *aadA10* which was within the integron and the beta-lactam gene *blaLCR-1* in the flanking region. Other studies have also shown that *qnrVC1* is a part of integrons that carry multiple resistance genes [22, 54]. The class 1 integron observed here had a typical structure containing promoter sequences and an attachment site (*attI*) required for integration and transcription of gene cassettes [57]. The *qnrVC1* gene in all four strains was associated with a perfectly matched gene recombination site (*attC*). The *attC* sequence of *qnrVC1* gene cassette had 100% identity with that of *Vibrio parahaemolyticus* (GenBank accession number EU436855), supporting the likelihood that the *Vibrionaceae* is a source of *qnr* in *P. aeruginosa* [58].

## Discussion

While fluoroquinolones are the preferred empirical therapy for corneal ulcers, which are often caused by *P. aeruginosa*, increasing resistance in this bacterium raises the concern about the efficacy of these antibiotics [4–11, 59, 60]. Acquired fluoroquinolone resistance genes are starting to be reported in clinical isolates of *P. aeruginosa* [20–22, 48]. However, the rate of carriage of acquired fluoroquinolone resistance genes in ocular isolates has not been reported to date. This study examined the prevalence of acquired fluoroquinolone resistance genes, *crpP* and *qnrVC1* genes, which have recently been shown to be quinolone-associated acquired resistance genes in *P. aeruginosa*. We used whole genomes of 33 strains isolated from corneal ulcers in the last 25 years isolates from Australia and India. Fluoroquinolone resistance was common in Indian isolates and mutations in QRDRs were associated with this increased MIC to all three fluoroquinolones tested. Possession of the acquired gene *qnrVC1* substantially increased the MICs whereas *crpP* was not associated with increased fluoroquinolone resistance in those isolates that carried *crpP* but lacked QRDRs mutations or *qnrVC1*. Both quinolone resistance genes were integrated into the chromosome and their associated genomic islands were broadly common between strains.

Fluoroquinolone resistance in *P. aeruginosa* is usually associated with mutations in the genes *mexR, nfxB*, and *mexT* that lead to overexpression of efflux pumps and QRDRs of *gyrA, gyrB, parC* and *parE* that alter drug target sites [1, 13–15]. In addition, horizontally transferred fluoroquinolone resistance genes such as *crpP* and *qnr*-variants have been recently reported in clinical strains of *P. aeruginosa* [20–22, 55]. Our BLAST analysis against the NCBI database showed that more than half of strains in the *P. aeruginosa* complete genome database contained homologues of *crpP* and this corresponded to the detection frequency of *crpP* in the ocular strains reported in this manuscript. *CrpP* carriage rate was higher in Indian isolates (74%) compared to Australian isolates (43%) and approximately half of the *crpP* carrying strains regardless of region of isolation were not resistant to any of the fluoroquinolones tested. Furthermore, fluoroquinolone resistant *crpP*+ strains also had target site mutations *gyrA* and *parC* and had acquired the quinolone resistance gene *qnrVC1*. Therefore, the role of *crpP* alone on fluoroquinolone resistance could not be ascertained for these strains. Given that *crpP* is ciprofloxacin specific and associated with low-level resistance, its presence may not necessarily be responsible for MICs that exceeded the clinical breakpoint [20].

The *crpP* gene was associated with similar genomic islands in all strains. These genomic islands were characterised by the possession of integrative conjugative elements and DNA replication factors. The *crpP* gene was originally reported in the plasmid pUM505 [51] but there are no other studies showing plasmid associated *crpP* in *P. aeruginosa*. Therefore, we performed the BLAST search against the NCBI plasmid database specific to *P. aeruginosa* and observed two matches (plasmids pKLC102 and pY89) out of 45 complete plasmids. Furthermore, the most recent ancestors of *crpP* positive strains appeared at least 50 years ago, which is earlier than the appearance of resistant subclades in the phylogeny. This result, together with finding of high divergence in *crpP* orthologues between strains and lower CAI compared to a highly expressed chromosomal gene (*rlpS*) [61] suggests that acquisition of the *crpP* gene was a relatively old evolutionary event in *P. aeruginosa*. In addition, the presence of CRISPR-Cas genes in several *crpP* positive strains supports the latter hypothesis and further suggests that *crpP* was integrated into the chromosome in an early event.

The plasmid-mediated quinolone resistance *qnr* gene was first reported in 1998 [16] and this corresponds with increased usage of fluoroquinolones in the 1980s [62]. Our analysis also revealed that resistant *P. aeruginosa* strains were phylogenetically separated from susceptible strains from the 1990s. The *qnr* genes protect DNA-gyrase and topoisomerase IV to prevent quinolone actions and confer low-level fluoroquinolones resistance, often below the clinical breakpoint [63, 64]. The *qnrVC1* gene is a mobile quinolone resistance gene that is associated with class I integrons [53]. *qnrVC1* and another variant *qnrVC6* have been reported from clinical *P. aeruginosa* strains isolated between 2007 and 2012 [21, 54]. However, in this study, the acquired fluoroquinolone resistance gene *qnrVC1* was observed in isolates sampled in 2017 and 2018, indicating that transferable fluoroquinolone resistance genes may be recently acquired or passed unnoticed in ocular isolates of *P. aeruginosa*. No strains contained *qnrVC6*. We also observed that *qnrVC1* was associated with a Tn*3* transposon, which has not been previously reported in *P. aeruginosa*. Integration of several other antibiotic resistance genes in these mobile genetic elements suggests these elements may concentrate antibiotic resistance genes. *QnrVC* is reported predominantly in water bacteria including *Aeromonas* spp., and *Acinetobacter* spp [65]. In this study the recombination site (*attC*) of all four *qnrVC1* alleles had 100% identity with aquatic bacteria *Vibrio parahaemolyticus*. This corresponds with our previous study in ocular isolates of *P. aeruginosa*, where acquired resistance genes closely matched with that of environmental isolates [52, 66].

However, the current study showed that *qnrVC1* carrying strains also had QRDRs mutations and were highly resistant to all three fluoroquinolones compared to those lacking *qnrVC1*. This indicates that possession of *qnrVC1* potentially facilitates selection of strains with high-level fluoroquinolone resistance. Given that *qnrVC1* has been reported as being responsible for low-level fluoroquinolone resistance [18], the high MICs observed for these isolates is not clear but may indicate some synergistic activity between the QRDR mutations and *qnrVC1*.

Like in many other studies, fluoroquinolone resistant strains were also associated with possession of *exoU*, which is the predominate virulent genotype of *P. aeruginosa* in ocular infections [67–77]. Although the reason behind this predominance is not completely understood, the concurrent occurrence of transferable fluoroquinolone resistance genes and *exoU* may heighten the concern of selection of more virulent strains during antibiotic therapy.

In conclusion, mobile genetic elements play a key role in the rapid spread of antibiotic resistance genes. This study provides evidence for carrying of fluoroquinolone resistance genes *qnrVC1* and *crpP* in ocular isolates of *P. aeruginosa*. The *qnrVC1* gene was mobilised by a class I integron and Tn3 transposon, and associated with other antibiotics resistance genes. The *crpP* gene may have evolved prior to other transferable fluoroquinolone resistance genes. Although possession of these genes has not been shown to be associated with high level of fluroquinolone resistance, *qnrVC1* in strains with *gyrA* and *parC* mutations could be associated with very high fluroquinolone resistance. Further studies on a larger number of fluoroquinolone resistance strains, including molecular analysis of combined effects of mutations in resistance, should be undertaken to provide further insights into the role of acquired fluoroquinolone resistance gene in clinically significant resistance in *P. aeruginosa*.

## Nucleotide accession

The nucleotide sequences are available in the GenBank under the Bio project accession number PRJNA590804 and PRJNA431326.

## Acknowledgements

The authors would like to acknowledge the Singapore Centre for Environmental Life Sciences Engineering (SCELSE), whose research is supported by the National Research Foundation Singapore, Ministry of Education, Nanyang Technological University and National University of Singapore, under its Research Centre of Excellence Programme. We are also thankful to UNSW high performance computing facility KATANA for providing us cluster time for data analysis.

