## Supplementary Information for "Acquired fluoroquinolone resistance genes in corneal isolates of *Pseudomonas aeruginosa*"

**Supplementary Table 1. Isolates of *Pseudomonas aeruginosa*.**

| ***P.aeruginosa* isolate designation** | **Year of isolation** | **Region of isolation** |
|---|---|---|
| PA17* | 1992 | Flinders University, Adelaide, Australia |
| PA40* | 1999 | SEH, Sydney, Australia |
| PA 176 | 2004 | Australia |
| PA182 | 2004 | PAH Brisbane Australia |
| PA149* | 2004 | Flinders University, Adelaide, Australia |
| PA123 | 2005 | PAH Brisbane Australia |
| PA126 | 2005 | PAH Brisbane Australia |
| PA127 | 2005 | PAH Brisbane Australia |
| PA 162 | 2006 | PAH Brisbane Australia |
| PA169 | 2006 | PAH Brisbane Australia |
| PA181 | 2006 | PAH Brisbane Australia |
| PA157* | 2006 | PAH Brisbane Australia |
| PA171* | 2006 | PAH Brisbane Australia |
| PA175* | 2006 | PAH Brisbane Australia |
| PA31* | 1997 | LVPEI, Hyderabad, India |
| PA32* | 1997 | LVPEI, Hyderabad, India |
| PA33* | 1997 | LVPEI, Hyderabad, India |
| PA34* | 1997 | LVPEI, Hyderabad, India |
| PA35* | 1997 | LVPEI, Hyderabad, India |
| PA37* | 1997 | LVPEI, Hyderabad, India |
| PA82* | 2004 | LVPEI, Hyderabad, India |
| PA188 | 2017 | LVPEI, Hyderabad, India |
| PA189 | 2017 | LVPEI, Hyderabad, India |
| PA193 | 2017 | LVPEI, Hyderabad, India |
| PA198 | 2017 | LVPEI, Hyderabad, India |
| PA202 | 2017 | LVPEI, Hyderabad, India |
| PA206 | 2017 | LVPEI, Hyderabad, India |
| PA216 | 2018 | LVPEI, Hyderabad, India |
| PA217 | 2018 | LVPEI, Hyderabad, India |
| PA218 | missing | LVPEI, Hyderabad, India |
| PA219 | 2018 | LVPEI, Hyderabad, India |
| PA220 | 2018 | LVPEI, Hyderabad, India |
| PA221 | 2018 | LVPEI, Hyderabad, India |

Isolates with asterisk (*) are from previous study (Subedi, Vijay, Kohli, Rice, & Willcox, 2018), LVPEI= LV Prasad Eye Institute, PAH=Princes Alexandra Hospital, SEH= Sydney Eye Hospital.

Subedi, D., Vijay, A. K., Kohli, G. S., Rice, S. A., & Willcox, M. (2018). Comparative genomics of clinical strains of *Pseudomonas aeruginosa* strains isolated from different geographic sites. *Sci Rep, 8*(1), 15668. doi:10.1038/s41598-018-34020-7

### Supplementary Figure 1: Tree view of the blast comparison of *qnrVC1* gene

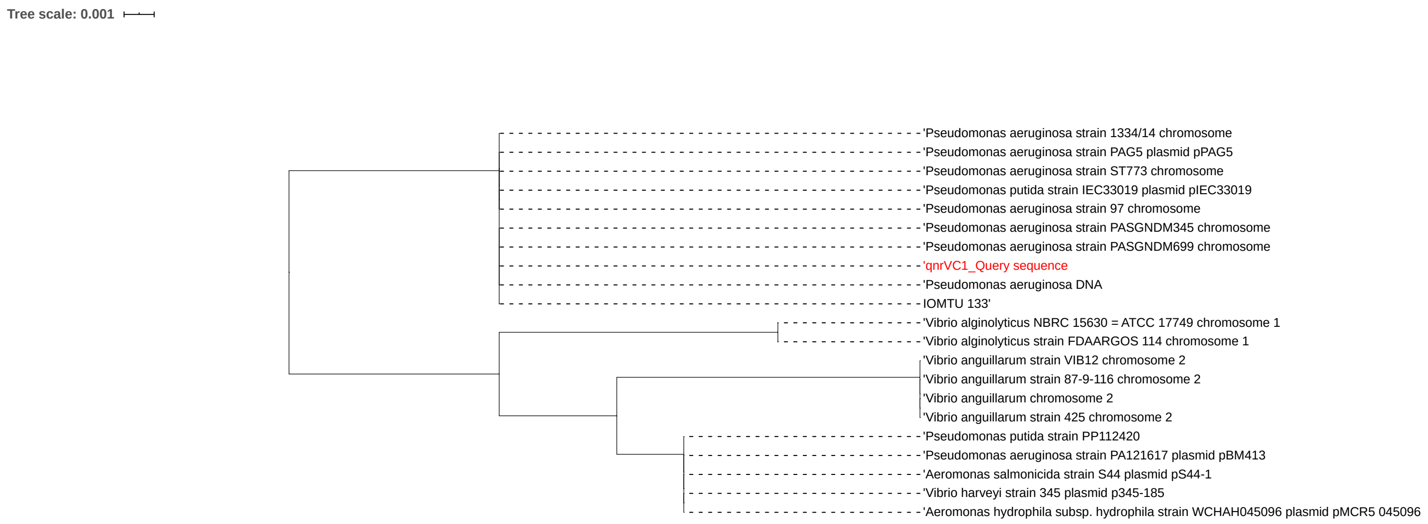
